# Relatively Shorter Muscle Lengths Increase the Metabolic Rate of Cyclic Force Production

**DOI:** 10.1101/2021.02.10.430661

**Authors:** Owen N. Beck, Jordyn N. Schroeder, Lindsey H. Trejo., Jason R. Franz, Gregory S. Sawicki

## Abstract

During animal locomotion, force-producing leg muscles are almost exclusively responsible for the whole-body’s metabolic energy expenditure. Animals can change the length of these leg muscles by altering body posture (*e.g.,* joint angles), kinetics (*e.g.,* body weight), or the structural properties of their biological tissues (*e.g.,* tendon stiffness). Currently, it is uncertain whether relative muscle fascicle operating length has a measurable effect on the metabolic energy expenditure of cyclic locomotion-like contractions. To address this uncertainty, we measured the metabolic energy expenditure of human participants as they cyclically produce two distinct ankle moments at three separate ankle angles (90°, 105°, 120°) on a fixed-position dynamometer exclusively using their soleus. Overall, increasing participant ankle angle from 90° to 120° (more plantar flexion) reduced minimum soleus fascicle length by 17% (both moment levels, p<0.001) and increased metabolic energy expenditure by an average of 208% (both p<0.001). Across both moment levels, the increased metabolic energy expenditure was not driven by greater fascicle positive mechanical work (higher moment level, p=0.591), fascicle force rate (both p≥0.235), or active muscle volume (both p≥0.122); but it was correlated with average relative soleus fascicle length (r=-179, p=0.002) and activation (r=0.51, p<0.001). Therefore, the metabolic energy expended during locomotion can likely be reduced by lengthening active muscles that operate on the ascending-limb of their force-length relationship.

## Introduction

Leg muscles govern walking and running performance. During the stance phase of walking and running, leg extensor muscles produce force in part to support and accelerate the body into the subsequent step. Concurrently, these force-producing leg muscles are almost exclusively responsible for the whole-body’s metabolic energy expenditure (Poole *et al.*, 1992; Griffin *et al.*, 2003; Marsh & Ellerby, 2006). Further, reducing an animal’s metabolic energy expenditure to perform a locomotor task increases how far they can travel at a given speed (Beck *et al.*, 2018) and how fast they can cover a fixed distance (Hoogkamer *et al.*, 2016). Thus, reducing the metabolic energy expenditure of an animal’s leg muscles while they continue fulfilling the physical requirements to sustain locomotion should improve locomotor performance.

Simply increasing the operating length of leg extensor muscles may decrease whole-body metabolic energy expenditure and improve locomotor performance. After all, the leg extensor muscles of many walking and running animals operate at shorter lengths than optimal (on the ascending limb of the force-length relationship) during ground contact (Roberts *et al.*, 1997; Biewener & Corning, 2001; Burkholder & Lieber, 2001; Daley & Biewener, 2003; Rubenson *et al.*, 2012; Bohm *et al.*, 2019). This is notable because muscles utilize more adenosine triphosphate (ATP) per unit of force production when they are maximally activated at shorter lengths than optimal (less economical force production) (Fig. 1) (Elzinga *et al.*, 1984; Stephenson *et al.*, 1989; Kentish & Stienen, 1994; Hilber *et al.*, 2001). Numerically, Hilber and colleagues (Hilber *et al.*, 2001) reported that compared to maximally producing force at an optimal length, sarcomere force production is 127% and 177% less economical at 0.8 and 0.6 of its optimal length, respectively. To envision the implications of these values during locomotion, the largest human ankle extensor muscle (soleus) is reported to produce force at 0.65 to 0.99 of its maximal voluntary contraction’s optimal length during the ground contact of walking and running (Rubenson *et al.*, 2012). These *in vivo* muscle lengths (*i.e.,* 0.65 to 0.99) may be overestimated due an inverse relationship between muscle activation and optimal fascicle length (Holt & Azizi, 2014; Hessel *et al.*, 2020). Furthermore, at shorter muscle lengths than optimal, active force production is reduced per unit of activation (Hill, 1953; Gordon *et al.*, 1966). Hence, for muscles to keep producing the force required to sustain locomotion, operating at shorter lengths than optimal requires the body to activate more muscle fibers and/or increase rate coding, both of which increase metabolic energy expenditure (Christie *et al.*, 2016; Beck *et al.*, 2019). Altogether, increasing the length of active leg muscles that operate at shorter lengths than optimal during stance likely decreases whole-body metabolic energy expenditure during locomotion.

**Figure 1.**
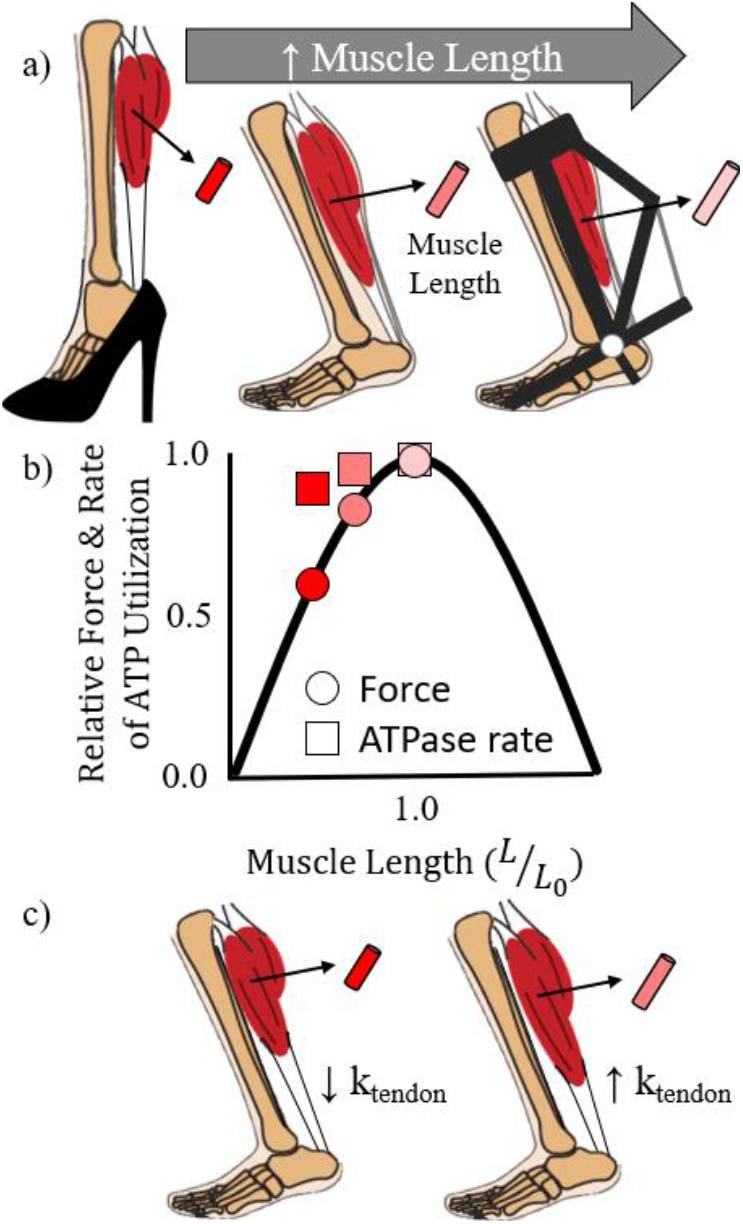
Illustrations of triceps surae fascicle lengths during the mid-stance of walking a) in high-heeled shoes (Csapo *et al.*, 2010; Cronin *et al.*, 2012), barefoot, with an ankle exoskeleton (Nuckols *et al.*, 2020), as well as c) barefoot with a more compliant and stiffer Achilles tendon (k_tendon_). b) Conceptual graph showing isometric muscle fascicle force production and adenosine triphosphate (ATP) utilization relative to optimal muscle operating length (*L*_0_) (Hilber *et al.*, 2001).

Despite the aforementioned rationale, muscle operating lengths are not often considered to have a *measurable* effect on whole-body metabolic energy expenditure during locomotion (Taylor, 1994; Minetti & Alexander, 1997; Pontzer, 2016; Kipp *et al.*, 2018; van der Zee & Kuo, 2020). This omission may be because the metabolic influence of producing force at different muscle lengths is typically studied during isometric contractions (Elzinga *et al.*, 1984; Stephenson *et al.*, 1989; Kentish & Stienen, 1994; Hilber *et al.*, 2001), rather than during cyclic length changing contractions that better mimic locomotion muscle mechanics. Another reason is that it is difficult to separate the metabolic effect of muscle operating lengths from other biomechanical parameters during locomotor-like contractions. For instance, when force-producing muscles shorten they perform mechanical work, and the further that a force-producing muscle shortens the more metabolic energy it expends (Fenn, 1924; Ortega *et al.*, 2015). Scientists commonly attribute this increased metabolic energy expenditure to greater muscle mechanical work (Fenn, 1924; Ortega *et al.*, 2015) rather than the muscle producing force at less economical lengths. While it is nearly impossible to experimentally disentangle the metabolic effect of muscle operating lengths from other metabolically-relevant biomechanical parameters during locomotion, such as force and work, a controlled experiment that emulates aspects of locomotion may be capable of accomplishing the task.

To help link walking and running biomechanics to metabolic energy expenditure, our goal was to determine the metabolic influence of cyclically producing force at different muscle fascicle lengths. To accomplish this goal, we quantified the fascicle mechanics and metabolic energy expenditure of human soleus muscles as they cyclically produced force at different relative lengths. Based on the notion that producing a given force at relatively shorter fascicle lengths increases metabolic energy expenditure (Elzinga *et al.*, 1984; Stephenson *et al.*, 1989; Kentish & Stienen, 1994; Hilber *et al.*, 2001; Beck *et al.*, 2019), we hypothesized that cyclically producing the same average force at relatively shorter fascicle lengths would increase metabolic energy expenditure.

## Methods

### Participants

Nine volunteers completed the protocol (average ± SD; age: 26.3 ± 2.6 years; standing height: 1.77 ± 0.07 m; mass: 74.9 ± 11.4 kg; resting metabolic power 87 ± 12 W; Achilles tendon moment arm during barefoot standing: 4.9 ± 0.4 cm; resting soleus fascicle length: 4.1 ± 0.6 cm; estimated maximum soleus fascicle shortening velocity: 182 ± 25 mm/s (Bohm *et al.*, 2019)). Prior to the study, each participant gave informed written consent in accordance with the Georgia Institute of Technology Central Institutional Review Board.

### Protocol

Participants arrived to the laboratory in the morning following an overnight fast. Upon arrival, participants laid supine on a dynamometer with custom attachments that supported their legs in the testing position: right knee and ankle supported at 50° and 90°, respectively (Fig. 2). 90° indicates perpendicular segments and more acute angles indicates joint (dorsi)flexion. In this position, participants rested for 10 minutes while breathing into a mouth piece that channeled expired air to a metabolic cart (TrueOne 2400, ParvoMedic, Sandy, UT, USA). Next, we shaved participant leg hair and used electrode preparation gel to lightly abrade the skin superficial to their right soleus, lateral gastrocnemius, and tibialis anterior (NuPrep, Weaver and Co., Aurora, CO). Subsequently, we placed a bipolar surface electrode over the skin superficial to each respective muscle belly and in the same orientation as the muscle fascicles (Delsys Inc., Natick, MA). We secured a linear-array B-mode ultrasound probe to the skin superficial of each participant’s right medial soleus (Telemed, Vilnius, Lituania). We placed reflective markers on the dynamometer at its axis of rotation, 10 cm above the axis of rotation, as well as on the participant’s skin/clothes superficial to their right leg’s medial knee-joint center, medial malleolus, and first metatarsal head (Fig. 2).

**Figure 2.**
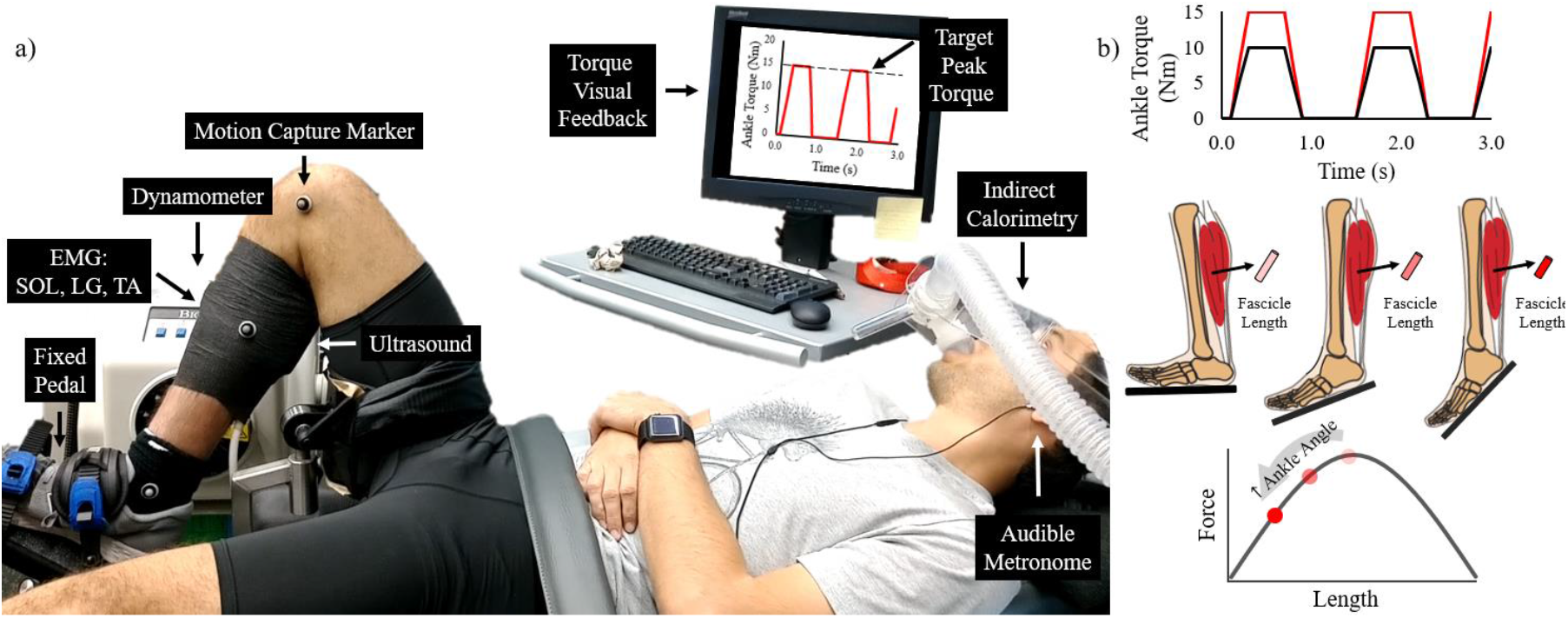
a) Experimental setup of a participant cyclically generating soleus muscle force to produce a plantar flexor moment that exerts an external torque on a fixed dynamometer pedal following the cues of an audible metronome and visual feedback. EMG, electromyography; SOL, soleus; LG, lateral gastrocnemius; TA, tibialis anterior. b) Illustrations of the two target torque levels (peak torque: 10 Nm and 15 Nm), three ankle angles (90°, 105°, and 120°) with the corresponding hypothetical minimum soleus fascicle operating lengths and their respective location on a muscle force-length relationship.

Next, participants performed four maximum voluntary contractions (MVCs) with their ankle joint center in-line with the dynamometer’s axis of rotation (Biodex Medical Systems Inc., USA): three plantar flexion MVCs and one dorsiflexion MVC. In a random order, participants conducted plantar flexor MVCs with their right knee at ~70°, 60° and 50° and their ankle at 90° (Rubenson *et al.*, 2012). Because MVC ankle moment did not increase with more extended knee angles, we deemed the contribution of the bi-articular gastrocnemius on ankle moment to be negligible (Rubenson *et al.*, 2012). Next, participants performed a dorsiflexion MVC with their right knee at 50° and ankle at 90° to maximally activate their tibialis anterior. At least two minutes of rest preceded each MVC to mitigate fatigue (Kawakami *et al.*, 2000).

Participants then performed six, five-minute trials with their knee at 50° separated by at least five minutes of rest. These trials consisted of each participant repeatedly producing plantar flexor moments on a fixed-position dynamometer foot-pedal at the downbeat of an audible metronome and then relaxing at the subsequent upbeat (metronome played at 1.5 Hz; Fig. 2). To guide ankle plantar flexor moments throughout each trial, participants watched a computer screen that displayed the trial’s target maximum dynamometer torque and the recorded dynamometer torque profile over the previous 5-10 s. Participants performed trials at each of the two dynamometer torque levels (10 Nm and 15 Nm) at the following ankle angles: 90°, 105°, and 120°. We randomized the trial order and collected rates of oxygen uptake and carbon dioxide production, dynamometer torque data (100 Hz), motion capture data (200 Hz) (Vicon Motion Systems, UK), soleus fascicle length and orientation (100 Hz), as well as the surface electromyography of the soleus, tibialis anterior, and lateral gastrocnemius (1000 Hz) (Fig. 2).

### Soleus fascicle mechanics

To determine soleus fascicle kinematics, we recorded B-mode ultrasound images containing the posterior-medial soleus. We recorded soleus fascicle images during 20 seconds in the last two minutes of the metabolic trials. Within these 20 s, we post-processed soleus fascicle lengths and pennation angles throughout six consecutive moment generation cycles using a semi-automated tracking software (Farris & Lichtwark, 2016). For semi-automated images that did not accurately track the respective soleus fascicle’s position, we manually redefined the desired fascicle. We filtered soleus fascicle angle and length using a fourth-order low-pass Butterworth filter (6 Hz) and took the derivative of fascicle length with respect to time to determine fascicle velocity.

To quantify soleus kinetics, we used a custom Matlab script (Mathworks Inc., Natick, MA, USA) that filtered motion capture data using a fourth-order low-pass Butterworth filter (6 Hz) and subtracted the resting dynamometer torque from the corresponding trial torque. We computed net dynamometer torque from 12 consecutive moment generation cycles that encompassed the analyzed fascicle kinematic data. Due to small fluctuations in dynamometer torque, we implemented a 1 Nm dynamometer torque threshold to decipher the duration of active force production. Using anthropometric measures and filtered data, we calculated the net ankle moment using dynamometer torque and the position of the ankle’s axis of rotation relative to the dynamometer’s axis of rotation, then we estimated the change in soleus muscle-tendon moment arm lengths at each ankle angle (Bobbert *et al.*, 1986). In turn, we divided net ankle moment (m_ank_) by the Achilles tendon moment arm length (r_AT_) to calculate muscle-tendon force. Next, we divided muscle-tendon force by the cosine of fascicle pennation angle (*θ*_p_) to calculate active soleus fascicle force (F_sol_).

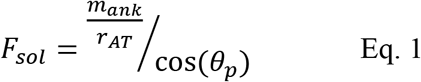

We assumed that passive muscle fascicle forces are negligible and we omitted the data from one five-minute metabolic trial because the participant achieved an average maximum ankle moment that was >5 Nm more than targeted. Further, we deemed optimal soleus fascicle length to be consistent across muscle activation magnitudes (de Brito Fontana & Herzog, 2016) and the same value that we measured during resting at a 90° ankle angle (Beck *et al.*, 2020). We set maximum fascicle shortening velocity to 4.4 resting lengths per second (Bohm *et al.*, 2019) based on the notion that only slow oxidative soleus fibers are active during sustained metabolic trials (Beck *et al.*, 2020).

### Biomechanical models

Recently, two studies performed similar experimental protocols and well-linked the mechanics of muscle fascicles cyclically producing force to metabolic energy expenditure. One study indicated that the combination of muscle fascicle force rate (*Ḟ*), positive mechanical work (*W*_+_), and force-time integral (∫ *Fdt*), scaled by corresponding cost coefficients (a, b, c), well-explains muscle metabolic energy expenditure (Eq. 2) (van der Zee & Kuo, 2020). The other study suggested that active muscle volume (*V*_act_) well-explains metabolic energy expenditure (*Ėmet*) during cyclic contractions that varied in duty factor (Beck *et al.*, 2020). Briefly, active muscle volume is calculated using active muscle fascicle force production (*F*_act_), optimal fascicle length (*l*_0_), stress (*σ*), and the fascicle’s force-length and force-velocity force potential (*FL* and *FV*, respectively) (Eq. 3) (Beck *et al.*, 2019). Due to the similarity of these previous studies to the present one, as a secondary objective of this paper we tested whether those muscle mechanics models could explain the present study’s metabolic data (Eq. 2 and 3).

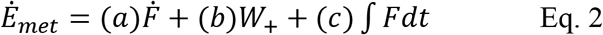

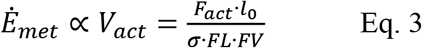

### Muscle activation

We band-pass filtered the raw soleus, lateral gastrocnemius, and tibialis anterior, electromyography signals between 20 and 450 Hz from the same 12 consecutive torque generation cycles that we used to assess net ankle moment. We full-wave rectified the filtered electromyography signals and calculated the root mean square of the rectified signals using a 40- millisecond moving window. Due to technical issues, we were unable to collect one participant’s tibialis anterior activation during the metabolic trials.

### Metabolic energy expenditure

During the resting trial and each cyclic force-production trial, we used open-circuit expired gas analysis to record the participant’s rates of oxygen uptake (V̇o_2_) and carbon dioxide production (V̇co_2_). We averaged V̇o_2_ and V̇co_2_ over the last minute of each trial and used a standard equation to calculate metabolic power (W) (Peronnet & Massicotte, 1991). Next, we subtracted each participant’s resting metabolic power from their experimental values to yield net metabolic power. We removed three metabolic data values (of 54) from our analyses because the corresponding respiratory exchange ratio did not reflect a respiratory quotient value that was indicative of fat and/or carbohydrate oxidation (Peronnet & Massicotte, 1991).

### Statistical analyses

Unless otherwise specified, we performed all statistical tests within the targeted lower and higher ankle moment trials independently. We performed t-tests to determine whether the targeted lower and higher cycle-average torque trials elicited different average ankle moments. We performed linear mixed models to determine the influence of ankle angle on the duration of active force production, force production cycle frequency, average ankle moment, average muscle-tendon force, average and maximum soleus fascicle pennation angle, average fascicle force, fascicle force-time integral, positive fascicle mechanical work, fascicle force rate, average and minimum fascicle operating lengths, maximal fascicle shortening velocity, average fascicle Hill-type force-length-velocity force potential, average soleus active muscle volume, soleus activation, lateral gastrocnemius activation, tibialis anterior activation, and net metabolic power. We also performed linear mixed models with two independent variables (average muscle fascicle length and positive mechanical work) and one dependent variable (net metabolic power). We performed independent linear regressions to determine the correlation between 1) average relative muscle fascicle length and 2) average soleus muscle activation on net metabolic power. We set the significance level (α = 0.05) and performed statistical analyses using RSTUDIO software (RSTUDIO, Inc., Boston, MA, USA).

## Results

Consistent with the study design, participants produced two distinct cycle average ± SD ankle moment levels: 4.85 ± 0.72 Nm and 6.58 ± 0.94 Nm (p<0.001) (Fig. 3). Within each moment level, the duration of active force production (both p≥0.158), force production cycle frequency (both p≥0.375), and cycle average ankle moment (both p≥0.678) each remained constant across ankle angles. However, not all metrics remained constant across ankle angles. Increasing ankle angle 30° lengthened participant Achilles tendon moment arms by 0.93 cm (Bobbert *et al.*, 1986), thereby decreasing average soleus muscle-tendon force (both p≤0.002) (Fig. 3). Greater ankle angles also increased average and maximum soleus fascicle pennation angles (both p≤0.001) (Fig. 3), which helped yield statistically similar cycle average soleus fascicle force production across ankle angles for the low ankle moment level (p=0.063) but not the higher ankle moment level (p=0.003) (Fig. 3). Similarly, at the lower ankle moment level, soleus fascicle force-time integral was independent of ankle angle (p=0.070), but it decreased by 19% due to increasing ankle angle from 90° to 120° within the higher moment level (p=0.003) (Fig. 4).

**Figure 3.**
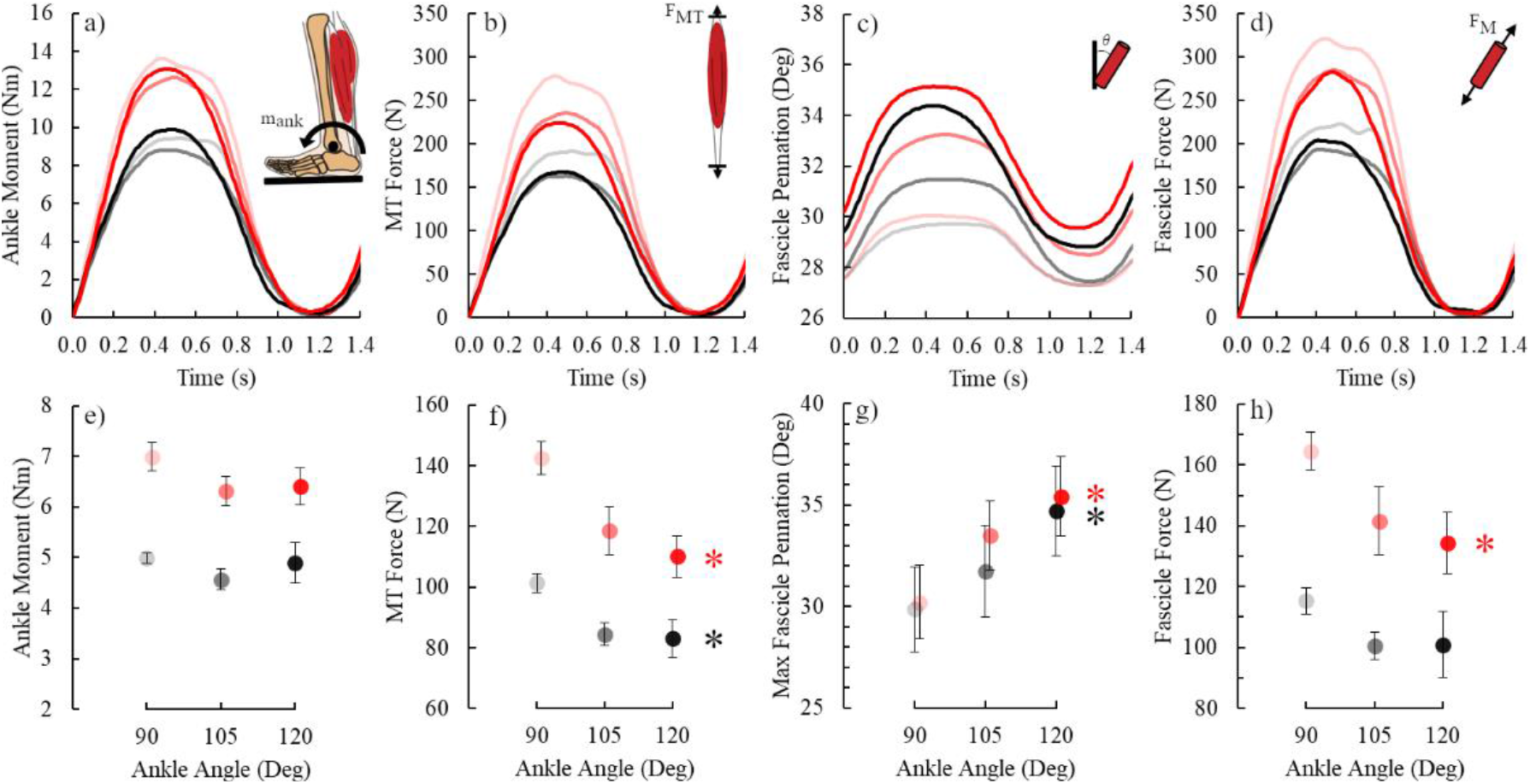
Top row: time-series plots of average a) ankle moment (m_ank_), b) muscle-tendon force (F_MT_), c) soleus fascicle pennation angle, and d) soleus fascicle force (F_M_). Bottom row: average ± SE e) average ankle moment, f) average MT force, g) maximum fascicle pennation angle, and h) average soleus fascicle force versus ankle angle. Black and red symbols are offset for clarity and indicate the lower and higher ankle moment levels, respectively. Lighter to darker colors indicate more dorsiflexed to plantar flexed ankle angles per moment level. Black and red asterisks (*) indicate that the corresponding moment level’s ankle angle affects the indicated dependent variable (p<0.05).

**Figure 4.**
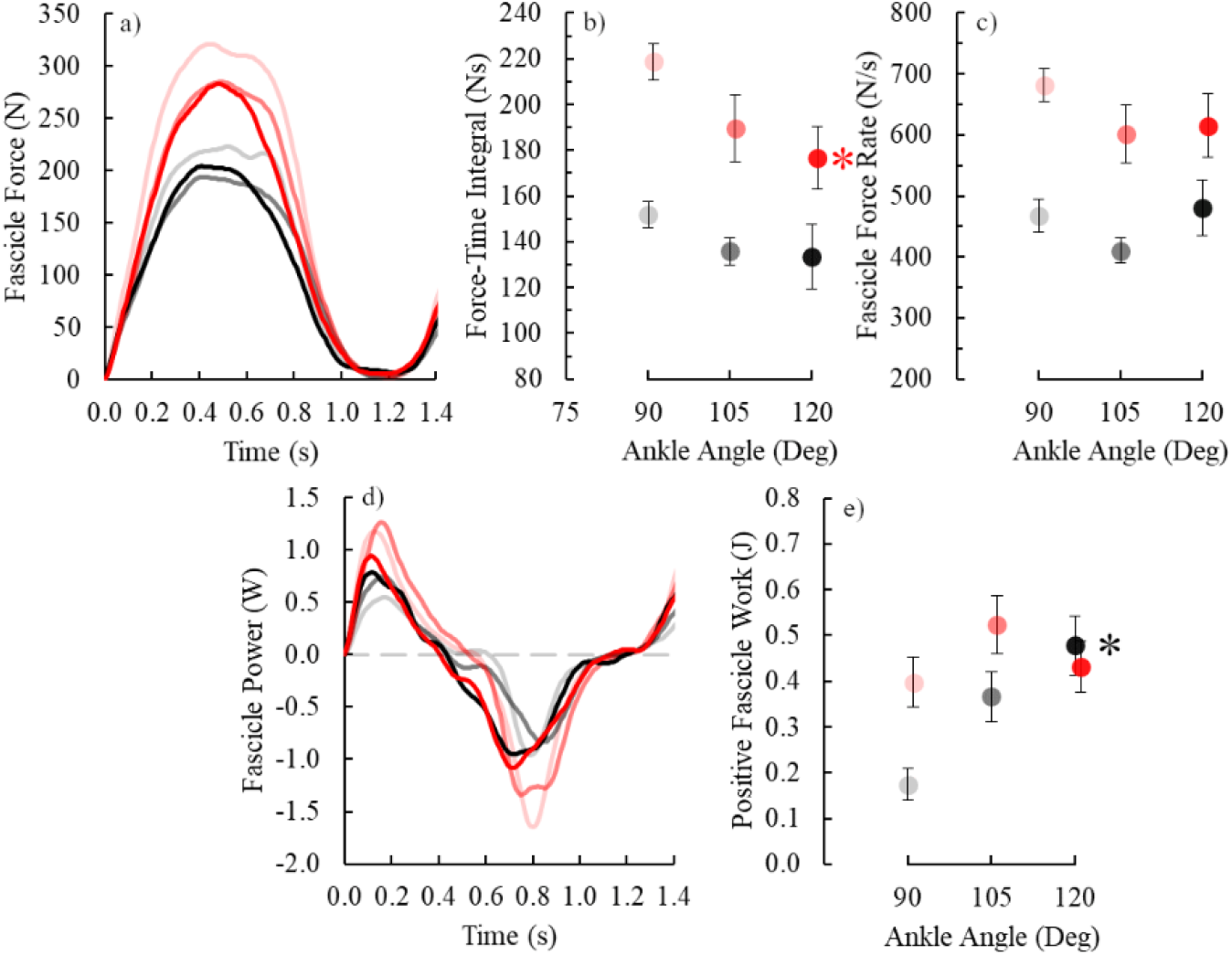
Time-series plots of average soleus fascicle a) force and d) power, as well as average ± SE soleus fascicle b) total force-time integral, c) force rate, d) and positive mechanical work. Black and red symbols are offset for clarity and indicate the lower and higher ankle moment levels, respectively. Lighter to darker colors indicate more dorsiflexed to plantar flexed ankle angles per moment level. Black and red asterisks (*) indicate that the corresponding moment level’s ankle angle affects the indicated dependent variable (p < 0.05).

Despite systematically shortening soleus fascicle lengths, increasing ankle angle did not alter many biomechanical parameters that previous dynamometer studies well-linked to net metabolic power. Specifically, both average and minimal soleus fascicle operating lengths decreased with increasing ankle angle (both p<0.001). These shorter fascicle operating lengths reduced the average soleus fascicle force-length potential by 7-8% across ankle moment levels (p<0.001) (Fig. 5). Further, greater ankle angles yielded faster maximum soleus fascicle shortening velocities at the lower ankle moment level (p<0.001), but not the higher ankle moment level (p=0.099). Combining cycle average fascicle force production and force-length-velocity potential (Beck *et al.*, 2019), ankle angle did not affect estimated cycle average soleus active muscle volume (both p≥0.122) (Fig. 5). Additionally, as aforementioned soleus fascicle force-time integral remained constant or slightly decreased with increased ankle angles (Fig. 3). Regarding the other metrics from equations 2 and 3 (Beck *et al.*, 2019; van der Zee & Kuo, 2020), soleus fascicle force-rate was independent of ankle angle (both p≥0.235) (Fig. 4), and positive soleus fascicle work increased across ankle angles within the lower ankle moment level (p<0.001), but not within the higher ankle moment level (p=0.591) (Fig. 4)..

**Figure 5.**
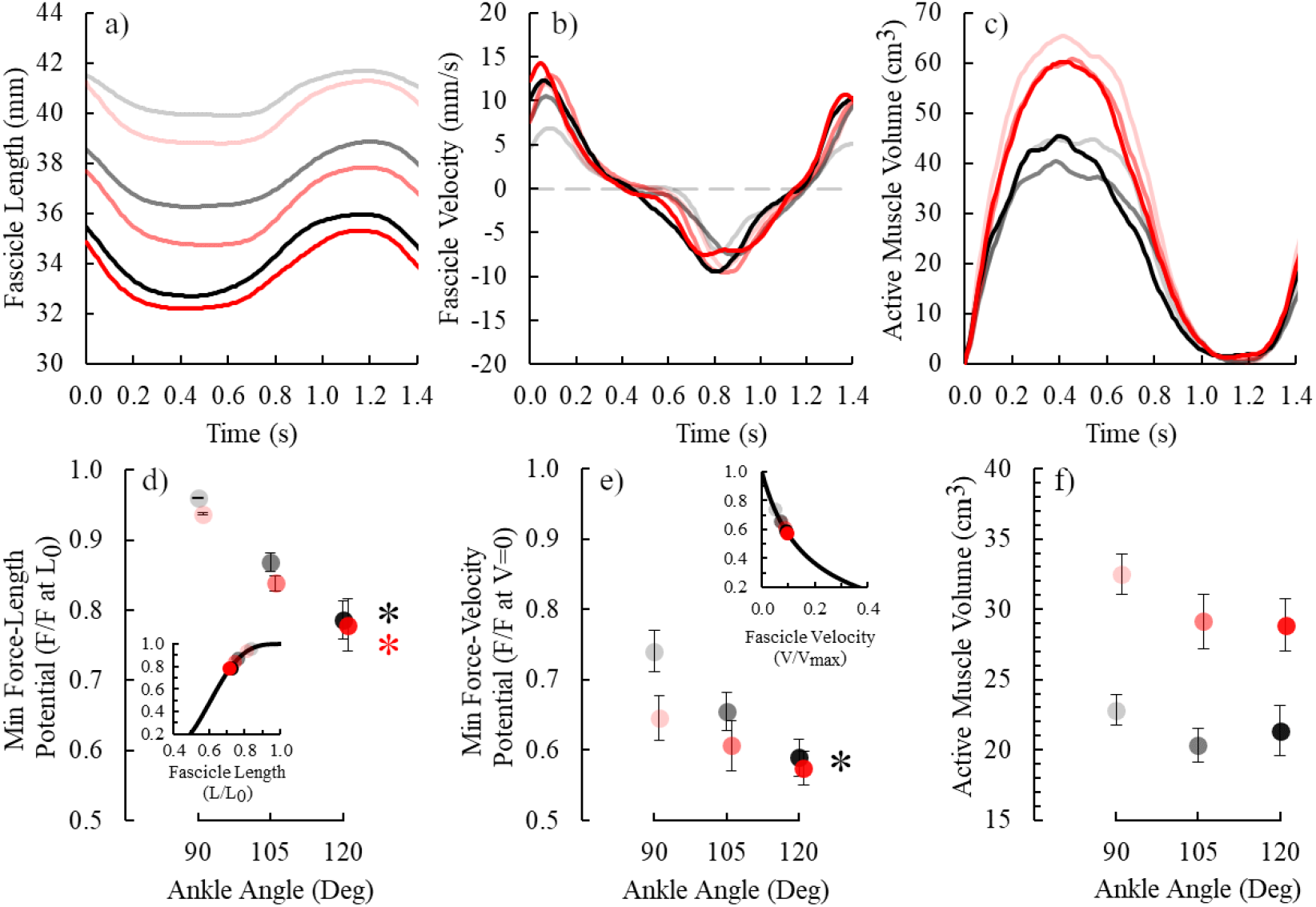
Top row: time-series plots of average soleus a) fascicle length, b) fascicle velocity, and c) active muscle volume. Bottom row: average ± SE d) minimum Hill-type force-length potential, e) minimum Hill-type force-velocity potential, and average f) active muscle volume versus ankle angle. Within panels d) and e) are the respective force-potentials plotted on the force-length and force-velocity curves, respectively. Black and red symbols are offset for clarity and indicate the lower and higher ankle moment levels, respectively. Lighter to darker colors indicate more dorsiflexed to plantar flexed ankle angles per moment level. Black and red asterisks (*) indicate that the corresponding moment level’s ankle angle affects the indicated dependent variable (p < 0.05).

Cyclically producing force at different ankle angles altered plantar flexor muscle activation. Both soleus and lateral gastrocnemius muscle activation increased by 140-200% with increasing ankle angle within each moment level (all p<0.001) (Fig. 6). Even though tibialis anterior activation statistically increased at greater ankle angles (both p≤0.027), we considered its influence on net metabolic power to be trivial because its cycle average activation was merely 2-5% of its MVC value across conditions.

**Figure 6.**
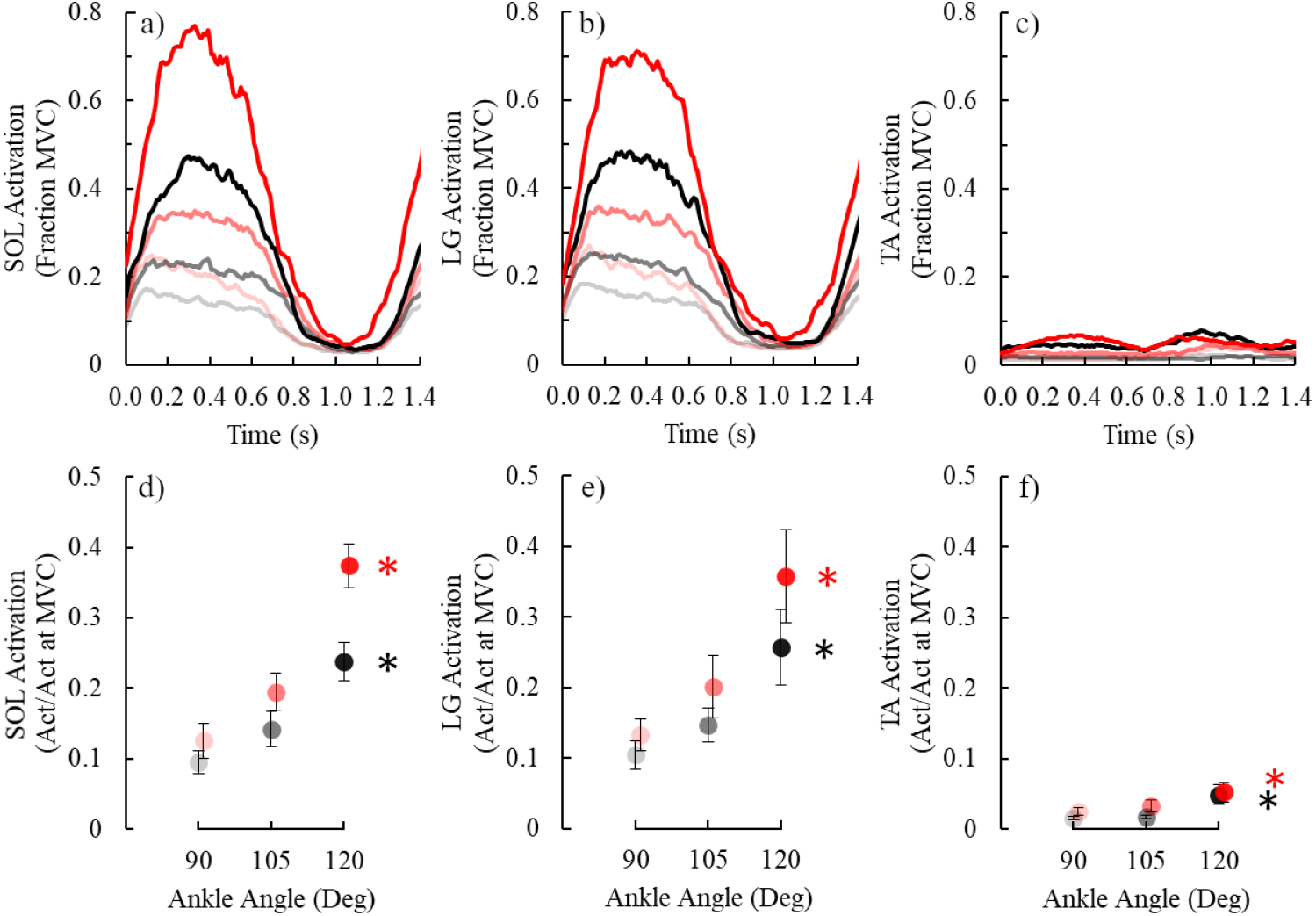
Top row: time-series plots of average a) soleus (SOL) activation (Act), b) lateral gastrocnemius (LG) activation, and c) tibialis anterior (TA) activation. Bottom row: average ± SE d) SOL activation, e) LG activation, and f) TA activation versus ankle angle. MVC is maximum voluntary contraction. Black and red symbols are offset for clarity and indicate the lower and higher ankle moment levels, respectively. Lighter to darker colors indicate more dorsiflexed to plantar flexed ankle angles per moment level. Black and red asterisks (*) indicate that the corresponding moment level’s ankle angle affects the indicated dependent variable (p<0.05).

Ankle angle affected the metabolic power of cyclic force production. Changing ankle angle from 90° to 120° increased net metabolic power by 189% and 228% within the lower and higher ankle moment levels, respectively (both p<0.001) (Fig. 7). Unlike previous dynamometer studies (Beck *et al.*, 2020; van der Zee & Kuo, 2020), neither the combined cost of muscle force-time integral, positive mechanical work, and force rate (Eq. 2); nor active muscle volume (Eq. 3) could explain the metabolic data (Suppl. Fig. 1). This is especially evident within the higher moment level where net metabolic power increased by 228% across ankle angles, but all of the following variables were either unchanged or decreased with increasing ankle angle (Suppl. Fig. 1): force-time integral (Fig. 4), force rate (Fig. 4), positive mechanical work (Fig. 4), and active muscle volume (Fig. 5). Further, within each moment level, positive mechanical work did not relate to net metabolic power while controlling for the influence of average fascicle length (p≥0.405). On the contrary, while controlling for positive mechanical work, decreasing average fascicle length was associated with an increased net metabolic power (both *β*=−1.4 to −3.1; p≤0.047). Pooled across ankle moment levels and participants, without controlling for other mechanical parameters, average relative muscle fascicle operating length inversely correlated with net metabolic power (linear regression: r=−179, p=0.002). Additionally, average soleus activation positively correlated with net metabolic power across ankle moment levels and participants (r=0.51, p<0.001) (Fig. 7).

**Figure 7.**
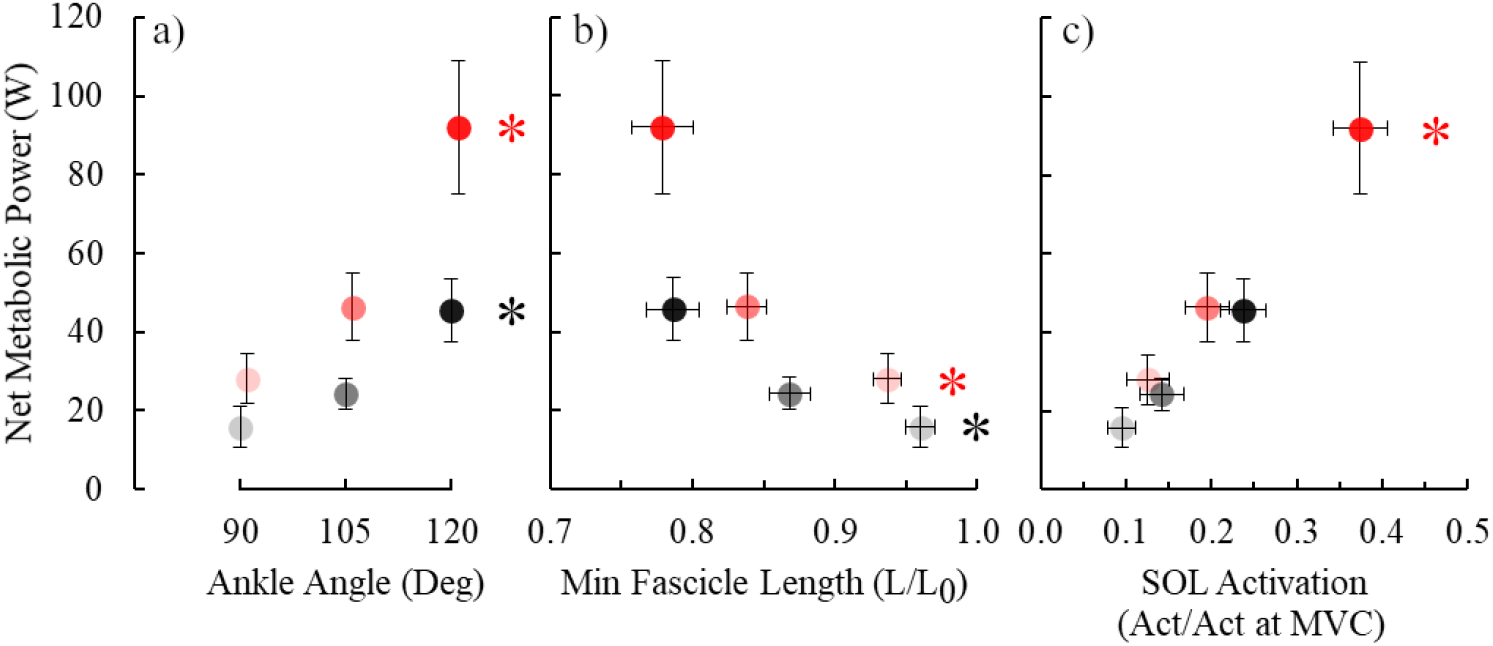
Average ± SE net metabolic power versus a) ankle angle, b) minimum fascicle length, and c) average soleus activation. Black and red symbols are offset for clarity and indicate the lower and higher ankle moment levels, respectively. Lighter to darker colors indicate more dorsiflexed to plantar flexed ankle angles per moment level. Black and red asterisks (*) indicate that the corresponding moment level’s ankle angle affects the indicated dependent variable (p < 0.05).

## Discussion

During locomotion, relative muscle fascicle operating lengths depend on the body’s posture, kinetics, and structural properties. In the present study, we controlled for participant limb-joint kinetics (constant cycle average ankle moment), structural properties (within participant design), and independently altered muscle fascicle operating lengths via postural changes (changing ankle angle). Using this protocol, we found that reduced muscle fascicle operating lengths *per se* increased metabolic energy expenditure during cyclic force production – supporting our hypothesis.

Why does cyclically producing force at relatively shorter muscle fascicle lengths increase metabolic energy expenditure? Muscle metabolic energy expenditure increases by either activating a greater volume of muscle or by expending more metabolic energy per unit active muscle volume (increased metabolic power density, W/cm^3^) (Beck *et al.*, 2019). Our results suggest that relative fascicle lengths primarily affect the latter. To explain, within the higher ankle moment level, changing ankle angle from 90° to 120° shortened average muscle fascicle operating lengths by 17% and decreased fascicle force-potential by 8% (Hill, 1953; Gordon *et al.*, 1966). To keep producing the same force profile, active muscle volume could theoretically increase by 8% (Hill, 1953; Gordon *et al.*, 1966). We would predict this change to elicit a mere 8% greater net metabolic power (Beck *et al.*, 2019), which is a far cry from the observed 228% increased metabolic energy expenditure. Therefore, even if there is a small increase in active muscle volume due to operating at shorter muscle fascicle lengths (*i.e.,* 8%), which we did not detect in our data, metabolic power density would need to increase by ~200% across ankle angles to match the net metabolic power data. In other words, producing force at relatively shorter fascicle lengths is likely metabolically expensive because muscles utilize ATP at faster rates, not because of the recruitment of more muscle fibers.

An increased metabolic power density from producing force at relatively shorter muscle lengths may be attributed to at least two factors. First, when sarcomeres produce force at shorter lengths than optimal, force production decreases faster than the corresponding ATP utilization (Fig. 1) (Elzinga *et al.*, 1984; Stephenson *et al.*, 1989; Kentish & Stienen, 1994; Hilber *et al.*, 2001). One proposed mechanism is that the net force produced by relatively shorter sarcomeres is mitigated by opposing forces that arise from compressed myofibrillar proteins. Another contributing factor that could help explain both greater muscle activation and net metabolic power at relatively shorter fascicle lengths is increased rate-coding (Enoka, 2015; Christie *et al.*, 2016). To continue producing a similar force magnitude at shorter fascicle lengths, participants may have increased the discharge rate of their soleus motor units, which would cycle ATP utilizing actin-myosin cross-bridges faster - thereby increasing metabolic power density.

It is important to recognize the difference between absolute and relative muscle fascicle operating length since they affect metabolic energy expenditure in different directions. Across vertebrates, the dimensions and quantity of actin-myosin cross-bridges per sarcomere are generally constant (Taylor, 1994; Burkholder & Lieber, 2001). This means that animals with optimally longer muscle fascicles typically have more sarcomeres in-series than animals with optimally shorter fascicles. Because whole-muscle force production depends on a muscle’s cross-sectional area and not its length, while considering other factors, absolutely longer muscle fascicles yield less economical force production due to more ATP consuming cross-bridges per unit force production (Taylor, 1994; Roberts *et al.*, 1998). However, shortening a muscle fascicle by a fixed distance (*e.g.,* 10 mm), shifts sarcomeres further down their ascending limb in an optimally shorter versus longer fascicle (*e.g.,* 40 mm versus 50 mm). Based on our data, decreasing a muscle fascicle’s length by a fixed distance likely increases metabolic energy expenditure more in an optimally shorter versus longer muscle fascicle. Hence, accounting for both absolute and relative muscle fascicle lengths may help explain the metabolic differences across participants with different optimal muscle fascicle lengths – such as in individuals pre- and post-injury (Williams & Goldspink, 1978; Hullfish *et al.*, 2019).

Accounting for both absolute and relative muscle fascicle lengths during locomotion may also help explain the differences in metabolic energy expenditure across animal species. For example, per unit body mass, smaller animals (*e.g.,* mice) have shorter muscle fascicles (Alexander *et al.*, 1981; Bennett, 1996) and expend more metabolic energy during locomotion than larger animals (*e.g.,* elephants) (Taylor *et al.*, 1970; Taylor *et al.*, 1982; Pontzer, 2007; Rubenson *et al.*, 2007). Based on the metabolic energy expenditure of producing force isometrically at an optimal fascicle length, the characteristically shorter muscle fascicles in smaller animals is conventionally viewed as an economical trait (Kram & Taylor, 1990; Roberts *et al.*, 1998; Kram, 2000). However, since many muscles in walking and running animals produce force while on the ascending limb of their force-length curve (Roberts *et al.*, 1997; Biewener & Corning, 2001; Burkholder & Lieber, 2001; Daley & Biewener, 2003; Rubenson *et al.*, 2012; Bohm *et al.*, 2019), optimally shorter muscle fascicle lengths may not necessarily translate to more economical locomotion. This may be especially true regarding comparisons across animals sizes, considering that smaller animals have relatively greater extensor muscle forces (Biewener, 1989) and more compliant tendons (Biewener, 2000) than larger animals; since both factors contribute to shorter muscle fascicles operating lengths. Hence, future studies that consider relative, in addition to absolute, muscle fascicle length during locomotion may better link walking and running biomechanics to metabolic energy expenditure across species.

There are many assumptions that may limit the findings of this study. First, consistent with previous studies (Beck *et al.*, 2020; van der Zee & Kuo, 2020), we assumed that the metabolic contribution of co-activating leg muscles was negligible. This assumption is based on the notion that there was likely slack in the biarticular gastrocnemius muscle-tendons (Rubenson *et al.*, 2012) and that the tibialis anterior muscle activation was trivial (Fig. 6). Second, we assumed that the soleus is primarily comprised of homogeneous muscle fibers (Johnson *et al.*, 1973), and that these fibers are exclusively recruited during the present study’s submaximal metabolic trials (Henneman, 1957). Hence, we deemed all active muscle fascicles in the soleus to have the same maximum shortening velocity, which we estimated to be 4.4 optimal lengths per second (Bohm *et al.*, 2019). As mentioned, we assumed that the soleus had uniform fascicle mechanics across the whole muscle, which is oversimplifies the complex architecture of the human soleus (Bolsterlee *et al.*, 2018). Nonetheless, passively changing muscle-tendon length alters soleus fascicle lengths and pennation angles change in the same direction across soleus muscle compartments (Bolsterlee *et al.*, 2018). Despite our many limitations and non-locomotor experiment, we find assurance when comparing our results to the most analogous locomotion experiment - walking in footwear with different heel heights. Similar to our study, increasing footwear heel height elicits postural changes that reduce triceps surae muscle fascicle forces (Simonsen *et al.*, 2012), relative muscle fascicle operating lengths (Csapo *et al.*, 2010; Cronin *et al.*, 2012), and increase whole-boy metabolic energy expenditure during walking and running compared baseline conditions (*i.e.,* barefoot or in flats) (Ebbeling *et al.*, 1994; Gu & Li, 2013).

In conclusion, our results suggest that operating further down the ascending limb of a muscle’s force-length curve may have a measurable influence on the metabolic energy expenditure during locomotion. Implications may help resolve why locomotion economy differs across and within animal species, in addition to informing biomechanical interventions that reduce user metabolic energy expenditure and consequently augment locomotor performance.

## Authors’ contributions

O.N.B. contributed to the conception and design of the study, acquisition of data, the analysis and interpretation of data, as well as the drafting of the article. J.N.S & L.H.T. contributed to acquisition of data and the revising of the article. J.R.F. contributed to the conception of the study, interpretation of data, as well as the drafting of the article. G.S.S. contributed to the conception and design of the study, the analysis and interpretation of data, as well as the drafting of the article. All authors approve of the manuscript and agree to be held accountable for all aspects of the work in ensuring that questions related to the accuracy or integrity of any part of the work are appropriately investigated and resolved.

## Data accessibility

Our data <u>will</u> be available on the corresponding and senior author’s personal websites upon acceptance (sites.google.com/view/owenbeck & sites.gatech.edu/hpl/publications/)

## Supplementary Figures

**Supplementary Figure 1.**
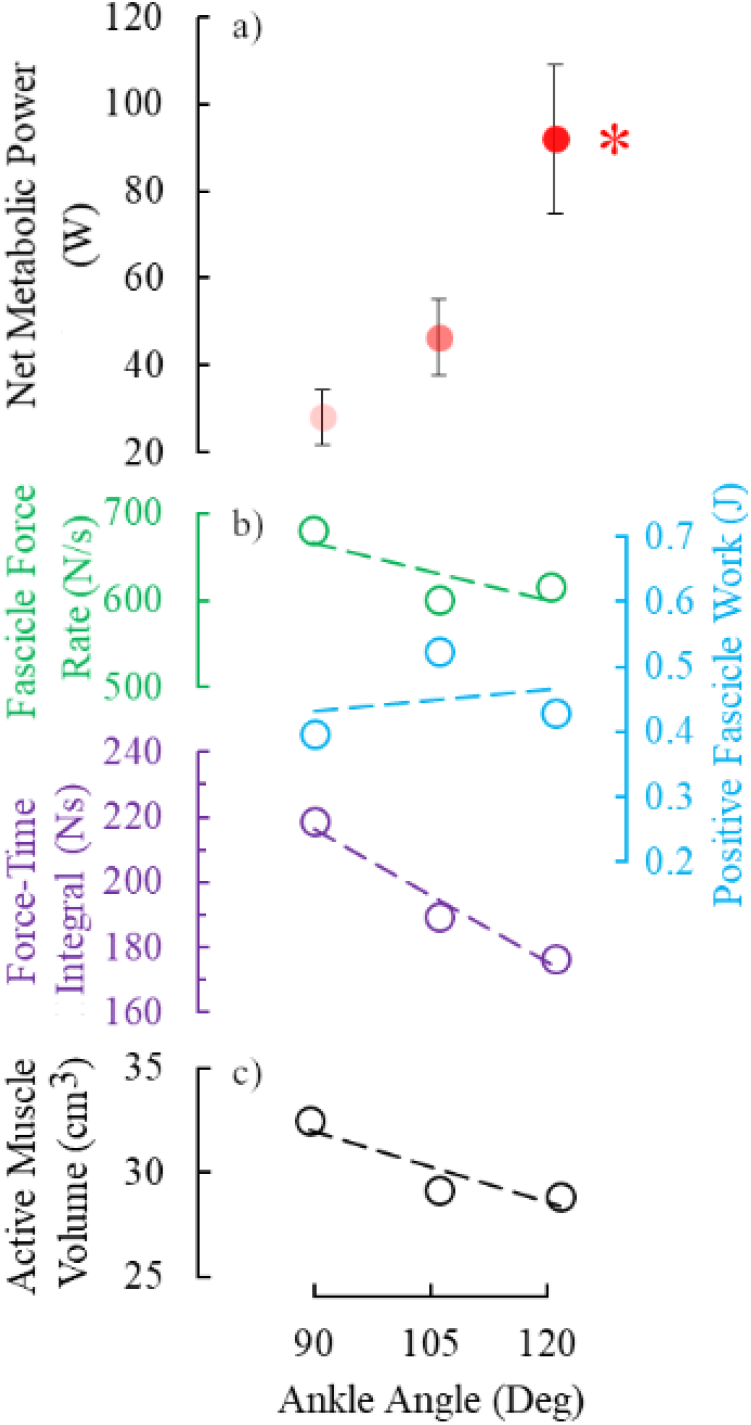
Net metabolic power is independent from the cost of 1) the sum of soleus fascicle force rate, positive fascicle work, and force-time integral, and 2) active muscle volume at the high ankle moment level (cycle average ankle moment: 6.57 Nm). a) Average ± SE net metabolic power versus ankle angle. b) Average soleus fascicle force rate, positive fascicle work, force-time integral, and c) active muscle volume versus ankle angle. Dashed lines indicate linear regression. For panels b and c, symbol color corresponds to the respective colored y-axis.

